# Darkness inhibits autokinase activity of bacterial bathy phytochromes

**DOI:** 10.1101/2023.12.15.571814

**Authors:** Christina Huber, Merle Strack, Isabel Schultheiß, Julia Pielage, Xenia Mechler, Justin Hornbogen, Rolf Diller, Nicole Frankenberg-Dinkel

**Affiliations:** Department of Microbiology, Rheinland-Pfälzische Technische Universität Kaiserslautern-Landau, 67663 Kaiserslautern, Germany; Department of Physics, Rheinland-Pfälzische Technische Universität Kaiserslautern-Landau, 67663 Kaiserslautern, Germany

**Keywords:** biliverdin, bathy phytochrome, dark sensor, far-red light sensor, histidine kinase, phosphorylation

## Abstract

Bathy phytochromes are a subclass of bacterial biliprotein photoreceptors that carry a biliverdin IXα chromophore. In contrast to prototypical phytochromes that adopt a Pr ground state, the Pfr-form is the thermally stable ground state of bathy phytochromes. Although the photobiology of bacterial phytochromes has been extensively studied since their discovery in the late 1990s, our understanding of the signal transduction process to the connected transmitter domains, which are often histidine kinases, remains insufficient. Initiated by the analysis of the bathy phytochrome *Pa*BphP from *Pseudomonas aeruginosa*, we performed a systematic analysis of five different bathy phytochromes with the aim to derive a general statement on the correlation of photostate and autokinase output. While all proteins adopt different Pr/Pfr-fractions in response to red, blue, and far-red light, only darkness leads to a pure or highly-enriched Pfr-form, directly correlated with the lowest level of autokinase activity. Using this information, we developed a method to quantitatively correlate the autokinase activity of phytochrome samples with well-defined stationary Pr/Pfr-fractions. We demonstrate that the off-state of the phytochromes is the Pfr-form and that different Pr/Pfr-fractions enable the organisms to fine-tune their kinase output in response to a certain light environment. Furthermore, the output response is regulated by the rate of dark reversion, which differs significantly from 5 seconds to 50 minutes half-life. Overall, our study indicates that bathy phytochromes function as sensors of light and darkness, rather than red and far-red light, as originally postulated.

## Introduction

Light is an essential environmental factor on earth. In addition to its function as a source of energy, actinic light can also represent a source of information in all domains of life (1). The ability to adapt behavior and physiology to ever-changing environmental conditions was increased during evolution through the presence of receptors that absorb light from a wide variety of spectral ranges. These photoreceptors are classified into different families depending on the specific wavelength by which they are activated, and the light absorbing chromophore that is embedded in the receptor: phytochromes, rhodopsins, xanthopsins, cryptochromes, LOV-(light-oxygen-voltage-sensing) domain-containing phototropins and BLUF-(blue-light sensing using FAD) domain proteins (2,3). Phytochromes are a class of red and far-red light sensing biliprotein photoreceptors (4). Originally, they were discovered in plants where they are involved in many developmental processes like seed germination, chloroplast development, and induction of flowering (5–7). The discovery in cyanobacteria, heterotrophic bacteria, and fungi occurred much later (8,9). Although the function of fungal or cyanobacterial phytochromes is partially understood (10–12), the role of phytochromes especially in heterotrophic bacteria is still enigmatic.

Nonetheless, the domain organization of bacterial phytochromes is largely conserved. The dimeric proteins consist of monomers containing an N-terminal photosensory core module (PCM) and a C-terminal regulatory output/transmitter module often representing a histidine kinase domain (HKD) (13–16). The amino-terminal PCM is composed of the Per-Arnt-Sim (PAS), followed by the cGMP-specific phosphodiesterase/adenylyl cyclase/FhlA (GAF) and the phytochrome-specific (PHY) domain (17–19). Together, the PAS and GAF domain form the chromophore-binding domain, harboring the linear tetrapyrrole chromophore biliverdin IXα (BV) (20,21). BV is autocatalytically attached via its A-ring to a conserved cysteine residue within the PAS domain establishing a thioether linkage with the C3^2^ atom. The chromophore is responsible for light reception and can adopt two (stable) protein conformational forms: the red light-absorbing Pr-form and the far-red light-absorbing Pfr-form (4,22). The corresponding conformational change is due to a reversible photoinduced *Z*-*E* isomerization of the C15=C16 double bond in the conjugated chromophore system between pyrrole ring C and D (Fig. S1) (18,23,24). The overall conformation of the tetrapyrrole chromophore in the Pr-form is *ZZZssa* and *ZZEssa* for the Pfr-form (25). In prototypical phytochromes, the Pr-form (λ_max_.∼700 nm, *Z* isomer) is the thermodynamically stable state and is dominant under dark conditions. In the dark, the Pfr-form (λ_max_.∼750 nm, *E* isomer) undergoes a thermally driven reversion (DR)) to the Pr-form (26).

Apart from prototypical phytochromes, the so-called bathy phytochromes have been described, thus far exclusively found in bacteria. Here, the far-red light-absorbing Pfr-form is the thermally stable, dark-adapted state of bathy phytochromes (2,27–29). The first bathy phytochrome discovered is derived from the human pathogen *Pseudomonas aeruginosa* (9,21). While bathy BphPs are widely distributed in rhizobial soil bacteria, like *Bradyrhizobium* (30), *Agrobacterium tumefaciens* (29), *Allorhizobium vitis* or *Rhodopseudomonas palustris* (31), they also exist in the plant pathogen *Xanthomonas campestris* (32) as well as in the Betaproteobacterium *Ramlibacter tataouinensis* (33), indicating that the distribution is not limited to the *Hyphomicrobiales* or to Alphaproteobacteria. Upon photoconversion from the Pr-to the Pfr-form, both, bathy and prototypical phytochromes undergo a structural change in the so-called ‘PHY tongue’ from a β-sheet structure to the formation of an α-helical structure (24,25,34). This tongue extends from the PHY domain to the chromophore-binding pocket in the GAF domain in kind of a hairpin structure (35). These structural changes lead to a reorientation of the GAF and PHY domain as well as the HKD which results in the modification of the dimer interface and thus confers the tongue a key role in signal transduction (17,36–39). Therefore, light reception results in changes in the absorption properties and the protein conformation, triggering autophosphorylation of the HKD on a conserved histidine residue. Subsequent phosphotransfer to a corresponding response regulator activates further signaling pathway within the cell (2).

Bacterial prototypical and bathy phytochromes have been extensively studied in terms of photobiology and three-dimensional structure. There is furthermore a fairly good understanding between phytochrome photostate and autokinase output in prototypical phytochromes with the highest kinase activity in the Pr-form (40–42). In contrast, there are only few reports on the relationship between photostate and autokinase output in bathy phytochromes (9,24,43). While our original work on the bathy phytochrome of *P. aeruginosa* (*Pa*BphP) demonstrated only marginal differences in autokinase output under different light conditions (9), Yang et al. and Mukherjee et al. reported a reduced kinase activity under dark conditions for *Pa*BphP (24,43). This ambiguity in experimental conditions and results does not allow for a clear correlation between the two parental photostates and the autokinase function. In this study, we provide a quantitative correlation between Pr/Pfr-fractions, calculated from UV/Vis spectra, and their corresponding biochemical autokinase readout. First established for *Pa*BphP, we apply this methodology systematically to a series of other bathy phytochromes from different Proteobacteria.

## Results

### Only darkness leads to a pronounced Pfr-form of *Pa*BphP

Although bathy phytochromes have been identified more than two decades ago, there is still a poor understanding of how signal input (light) is correlated with signal transmission (often autokinase activity). The cause of this lies in the consistent observation of a mixture of Pr- and Pfr-forms by numerous researchers when exposing the sample to red light, creating a challenge in establishing a clear correlation to autokinase output. Therefore, we decided to reinvestigate the photostates of the model bathy BphP of *P. aeruginosa* (*Pa*BphP) under various light conditions, including blue light and darkness. In order to isolate *Pa*BphP under most native conditions, the *bphP* gene, encoding the full-length phytochrome, was homologously expressed within *P. aeruginosa.* The purification was performed under additional supply of the BV IXα chromophore in the bacterial lysate.

Illumination of the purified protein with red (667 nm) or blue (426 nm) light (Fig. S4) resulted in thermally equilibrated Pfr-enriched forms of the phytochrome, exhibiting a peak at 700 nm and an absorption shoulder at 750 nm. As previously described (9,24), illumination with far-red light (791 nm) converted *Pa*BphP into a distinct and pure Pr-form with a peak at 700 nm, lacking the 750 nm shoulder (Fig. 1A; Fig. S2). Only prolonged incubation in darkness (> 6 h) yielded a pure Pfr-form. Employing these pure form absorption spectra of Pr and Pfr, we developed a calculation method to determine the fractions of the two conformers in a given sample (cf. experimental procedures, Fig. S6, Table S1). It turns out that, at the given wavelengths, red light illumination leads to approximately equal amounts of Pr- and Pfr-forms, blue light illumination slightly shifts the ratio towards the Pr-form (67/33 (Pr/Pfr)). Note that only far-red light and darkness are able to lead to fully developed 100 % Pr and Pfr-forms, respectively (Tab. 1).

**Fig. 1:**
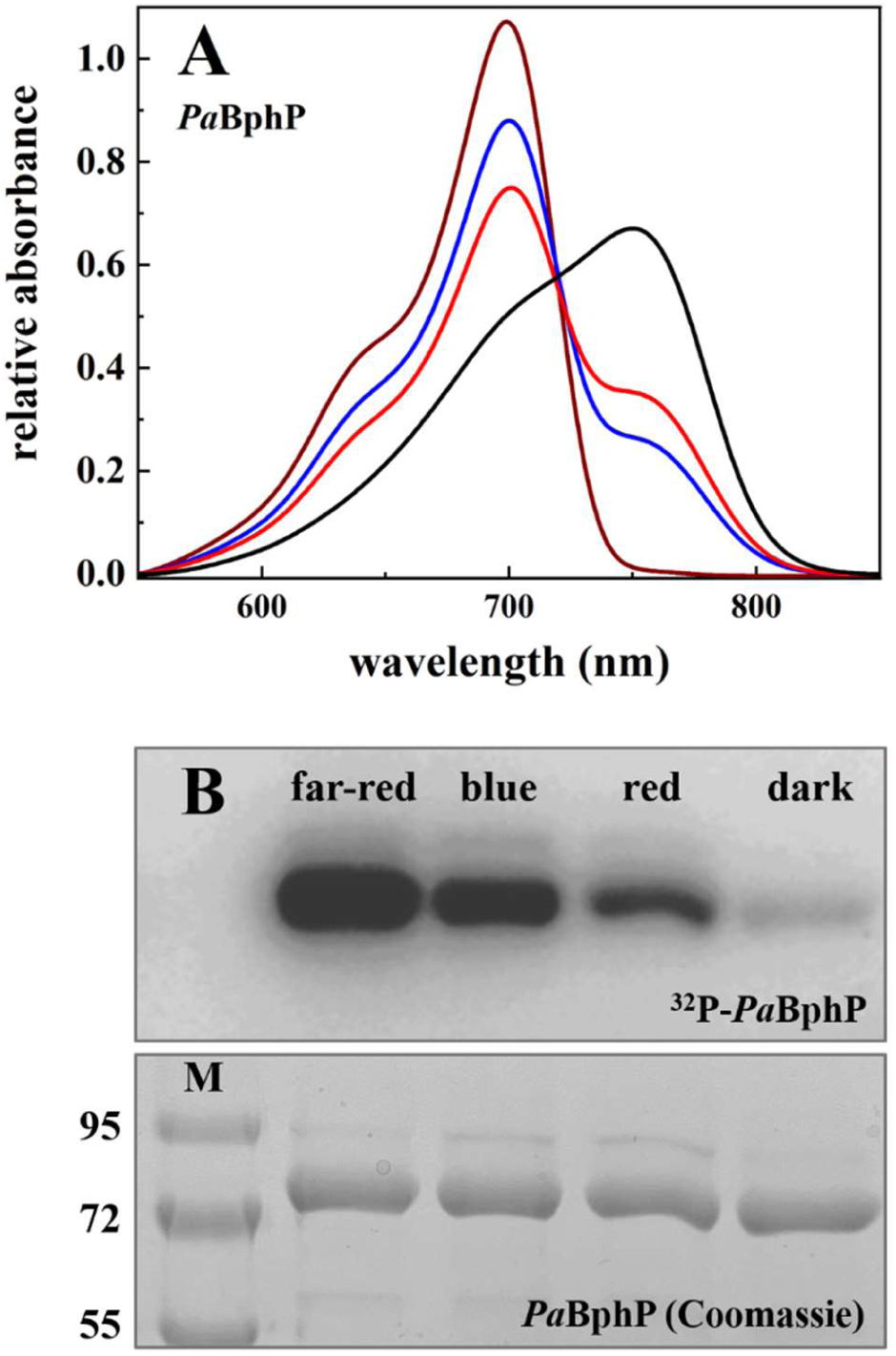
Spectral absorbance properties and autophosphorylation activity of recombinant produced *Pa*BphP under different light conditions. **A**. Absorbance spectra of *Pa*BphP (mathematically smoothed with Origin, Version 2022. OriginLab Corporation, Northampton, MA, USA. “Loess, Use Proportion for Span, Span 0.1”) after illumination with far-red light (791 nm, 5 min, far-red line), with blue light (426 nm, 10 min, blue line), with red light (667 nm, 10 min, red line), and after dark reversion (24 h, black line). The extend of conversion cannot be improved by longer exposure times. All data are generated from a single set of measurements. BV IXα was added in excess after cell lysis before purification. The exact ratios for each spectrum are also shown in Tab. 1. The errors for α amount to ±1% for Pfr and ±8% for Pr. **B.** Autoradiogram (top) and corresponding SDS-PAGE gel (10 % (w/v), bottom) after *in vitro* radiolabeling of *Pa*BphP with [γ-^32^P]-ATP under far-red (791 nm), blue (426 nm), and red light (667 nm) as well as under dark conditions. Autophosphorylation reaction was terminated after 5 min and the protein samples were separated by gel electrophoresis. Proteins were pre-exposed to the appropriate light for 5 min or incubated dark overnight before adding the radiolabeled ATP. *Pa*BphP has a relative molecular weight of approximately 83 kDa (M = marker; Color Prestained Protein Standard, Broad Range, NEB).

**Table 1:** Different Pr/Pfr-fractions for recombinant purified phytochromes of *P. aeruginosa*, *A. tumefaciens*, *A. vitis*, *R. tataouinensis*, *X. campestris,* and various variants of the proteins under dark, red, blue, or far-red light conditions employed for kinase assays shown in Fig. 3. The errors amount to ±1% for Pfr and ±8% for Pr.

|  | Pfr (%) | Pr (%) | Pfr (%) | Pr (%) | Pfr (%) | Pr (%) | Pfr (%) | Pr (%) |
| --- | --- | --- | --- | --- | --- | --- | --- | --- |
|  | dark |  | red light (667 nm) |  | blue light (426 nm) |  | far-red light (791 nm) |  |
| **PaBphP** | 100 | 0 | 50 | 50 | 33 | 67 | 0 | 100 |
| **AtBphP2** | 100 | 0 | 66 | 34 | 81 | 19 | 4 | 96 |
| **AtBphP2_D783N** | 97 | 3 | 57 | 43 | 72 | 28 | 0 | 100 |
| **AvBphP2** | 97 | 3 | 81 | 19 | 56 | 44 | 1 | 99 |
| **AvBphP2_D793A** | 92 | 8 | 78 | 22 | 60 | 40 | 0 | 100 |
| **RtBphP2** | 100 | 0 | 53 | 47 | 83 | 17 | 2 | 98 |
| **XccBphP** | 72 | 28 | 56 | 44 | 13 | 87 | 0 | 100 |
| **XccBphPΔPAS9** | 96 | 4 | 57 | 43 | 16 | 84 | 0 | 100 |

### The Pr/Pfr-fractions influence *Pa*BphP autokinase activity

Next, we investigated whether there are variations in the autokinase activity of the different illuminated samples. To accomplish this, a special illumination device was developed, allowing for light exposure during the radioactive kinase assays, comparable to that of the UV/Vis spectroscopy (Fig. S5). This was absolutely essential for kinase assays in the dark and samples with fast dark reversion rates. The assay revealed that the phytochrome autophosphorylates under all light conditions, but the activity is strongly inhibited and can be considered to be turned off in darkness (Fig. 1B). Depending on the wavelength of the incident light and thus the resulting protein conformation, *Pa*BphP displays different levels of phosphorylation. Autokinase signals increased – at the given wavelength - from red to blue to far-red, according to the correspondingly increasing Pr-fraction and decreasing Pfr-fraction (Table 1). Although there is a strong correlation between autokinase readout and the Pr-fraction in this specific case, the signal intensities observed in an autoradiogram cannot be reliably used to quantify kinase activity as a whole. This limitation is due to their dependence on exposure time and other experimental variables like the specific activity of the radionuclide. For a more accurate analysis and correlation of phytochrome activity, we propose to use the Pr/Pfr-fractions calculated from the spectroscopic data assuming that only the Pr-proportion of a given sample contributes to autokinase activity of the phytochrome. For the first time, we are able to determine the percentage of the active kinase in a given sample using this novel method.

To further validate the method, the autophosphorylation behavior during dark reversion of *Pa*BphP was tested (Fig. 2). After 5 min of far-red irradiation, resulting in a pure Pr-form and highest autokinase activity, the sample was transferred to complete darkness. In the dark, the formation of Pfr proceeds over time (Fig. 2A), correlating with a reduction in kinase activity as the proportion of Pr decreases (Fig. 2B, C). After 120 min, the sample still contains 13 % of Pr which is responsible for the weak kinase activity signal. Only after six hours in darkness, the sample is completely in the Pfr-form resulting in no kinase activity (see also Fig. 1B).

**Fig. 2.**
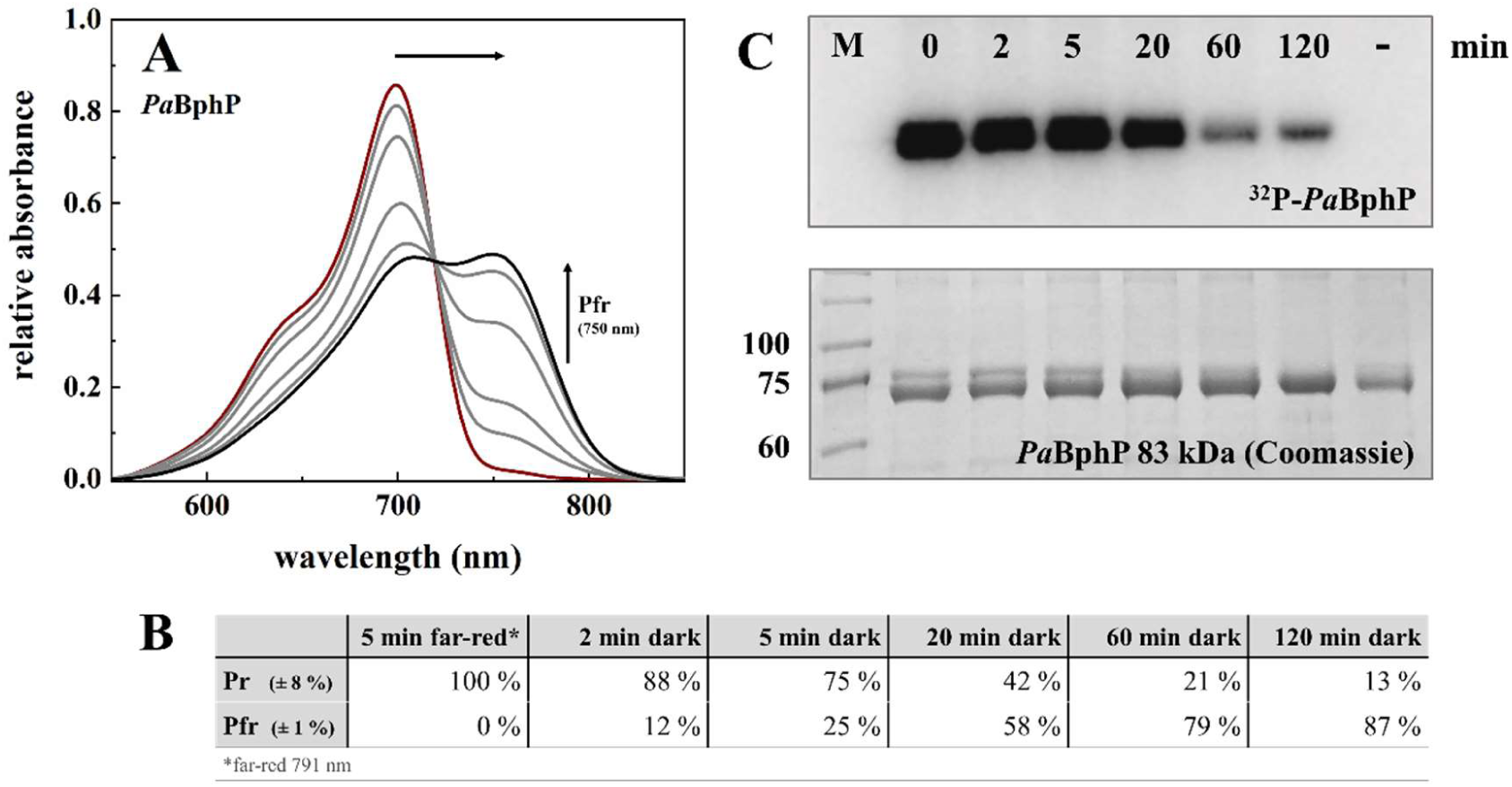
Correlation between time resolved dark reversion and autokinase activity of *Pa*BphP. **A.** Absorption spectra of *Pa*BphP (mathematically smoothed using Origin, Version 2022. OriginLab Corporation, Northampton, MA, USA. “Loess, Use Proportion for Span, Span 0.1”) after irradiation with far-red light (791 nm, 5 min, dark red line) and different time points during dark reversion (2, 5, 20 and 60 min gray lines; 120 min black line). Course of the dark reversion (increase of Pfr-form, decrease of Pr-form) is indicated by the arrows. All data are generated from a single set of measurements. BV IXα was added in excess after cell lysis prior to purification. **B.** Absorption of far-red light results in a distinct Pr-form, and dark incubation results in increasing formation of Pfr-form. The exact fractions of Pr and Pfr for each spectrum are given in the table. The errors for α amount to ±1% for Pfr and ±8% for Pr. **C.** Autoradiogram (top) and corresponding Coomassie-stained SDS-PAGE gel (10 % (w/v); bottom) after *in vitro* radiolabeling of *Pa*BphP with [γ-32P]-ATP under far-red (791 nm) light conditions (t = 0 min) and during the dark reversion after 2, 5, 20, 60 and 120 min. The minus sign (-) indicates the negative control *Pa*BphP_H513A, which displays no autophosphorylation signal after 120 min incubation with radiolabeled ATP. *Pa*BphP has a relative molecular weight of approximately 83 kDa (M= marker: Prestained protein marker, proteintech).

### All bathy phytochromes share a very pronounced Pr-form under far-red illumination and strongly enriched Pfr-forms in darkness

Since published data often use slightly different light conditions and are therefore difficult to compare, we started a comprehensive investigation of various bathy phytochromes from Proteobacteria to answer whether they display similar absorption properties under identical light settings. Although they all share the domain organization of the PCM, their output/transmitter domains differ (Fig. S3). While BphP2 of *R. tataouinensis* completely shares its domain organization with *Pa*BphP, those BphPs from *Agrobacterium* and *Allorhizobium* possess an HKD subfamily, a so-called HWE kinase domain in addition to a C-terminal response regulator domain. BphP of *Xanthomonas* completely lacks a kinase output domain but carries an PAS9 domain, likely involved in protein-protein interaction instead (44).

Far-red irradiation (791 nm) led to the photoconversion of all tested phytochromes into a pure Pr-species. The far-red light adapted form of *Xcc*BphP and *Xcc*BphPΔPAS9 is thereby most similar to the Pr-form of *Pa*BphP with no absorption at 750 nm (Fig. 3A-C; Tab. 1). However, the Pr λ_max_ of the *X. campestris* phytochrome is shifted to a shorter wavelength (686 nm) in comparison to *Pa*BphP (Tab. 2). For the two bathy phytochromes from *Agrobacterium* and *Allorhizobium*, protein variants, lacking the phosphor-accepting aspartate within the receiver domain, were employed. Previous work indicated that this amino acid residue significantly influences kinase activity but not the photocycle (42). Illumination of *At*BphP2_D783N and *Av*BphP2_D793A with far-red light resulted in two very similar spectral forms with minimal absorption at 750 nm (Fig. 3D, E) and Pr λ_max_ of 701 and 698 nm (Tab. 2), respectively. The Pr-form of *Rt*BphP2 displayed a very broad spectrum with an unusually low amplitude (Fig. 3F) - Pr λ_max_ is at 684 nm (Tab. 2). Based on the spectra of the respective pure forms of Pr and Pfr, for each phytochrome the Pr/Pfr- fractions were determined as described for *Pa*BphP (Tab. 1). Contrary to the pronounced Pr-form under far-red conditions, illumination with red and blue light always resulted in a mixture of Pr- and Pfr-forms (Tab. 1). Observed variations in Pr/Pfr-fractions in different phytochromes yet under the same illumination conditions are attributed to phytochrome-specific parameters of the respective photochemical equilibrium, i.e. λ_max_ of Pr and Pfr in the Soret- and Q-band region (Fig.S2 and Tab. 2), corresponding extinctions coefficients and photoisomerization quantum yields as well as rates for dark reversion. Only incubation in darkness led to the highest amount of Pfr-form in all bathy phytochromes. This is a major difference to the prototypical phytochromes where the highest amount of Pfr is only produced in red light. The accumulation of the Pfr-form in darkness was observed for all phytochromes studied and represents the characteristic of bathy phytochromes. However, *Xcc*BphP and *Av*BphP2_D793A accumulated hardly any pure Pfr state (Fig. 3B, E). This is in slight contrast to the published spectra of *Av*BphP2 that show a pure Pfr-form when incubated in the dark (31) but might be due to different experimental set-ups in the present study. For *Xcc*BphP, the truncated version without the PAS9 output domain performed as a typical bathy phytochrome and displayed a stable state in the homogenous Pfr-form (Fig. 3C). This is likely due to interference of the PAS9 domain with chromophore binding and the thermodynamics of dark reversion (32).

**Fig. 3:**
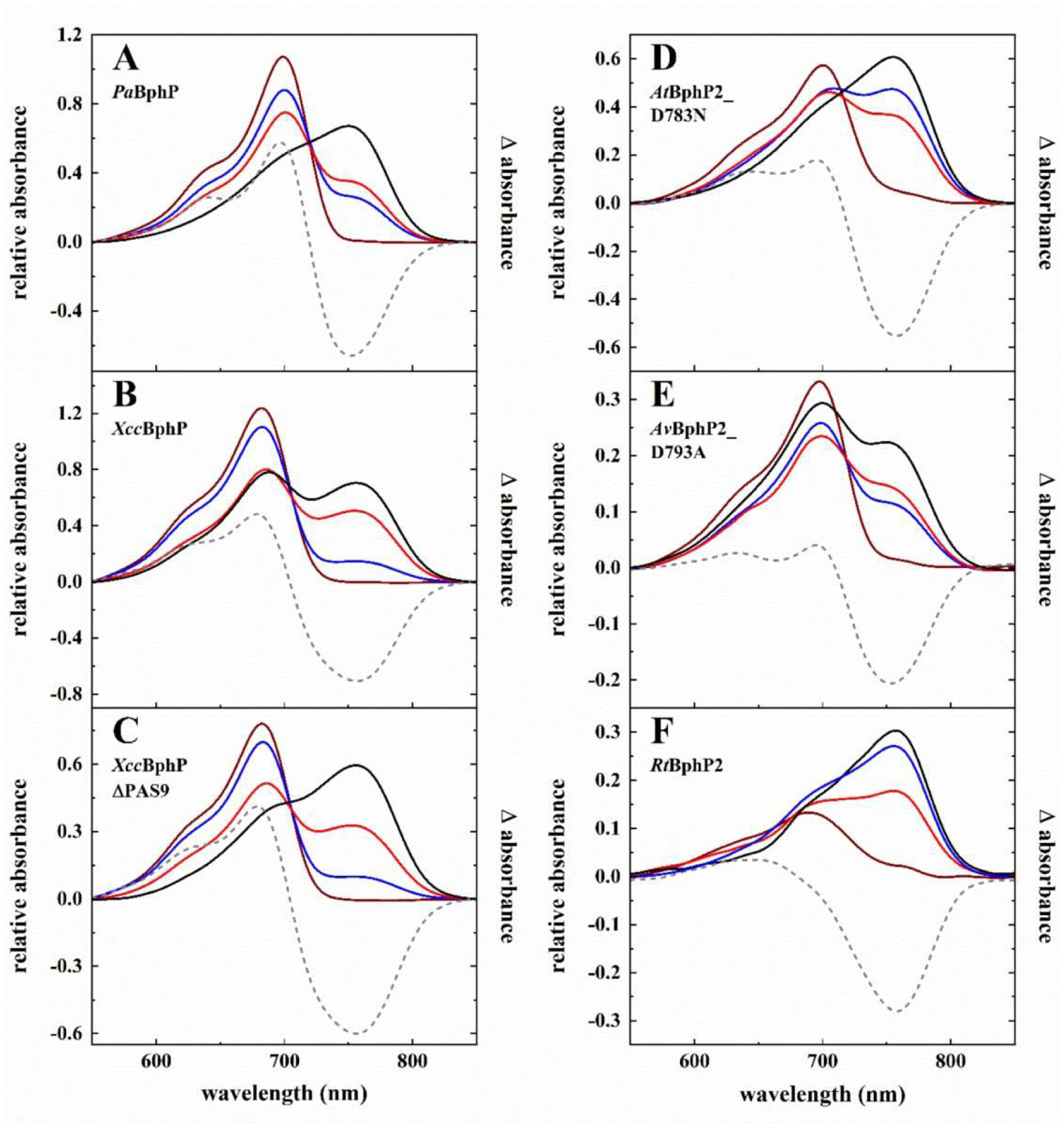
Spectral absorption properties of recombinant produced *Pa*BphP, *Xcc*BphP, *Xcc*BphPΔPAS9, *At*BphP2_D783N, *Av*BphP2_D793A and *Rt*BphP2. Absorbance spectra (mathematically smoothed with OriginLab, Version 2022. OriginLab Corporation, Northampton, MA, USA. “Loess, Use Proportion for Span, Span 0.1”) of *Pa*BphP **(A)**, *Xcc*BphP **(B)**, *Xcc*BphPΔPAS9 **(C)**, *At*BphP2_D783N **(D)**, *Av*BphP2_D793A **(E)** and *Rt*BphP2 **(F)** after illumination with far-red light (791 nm, 5 min, far-red lines), with blue light (426 nm, 5 min B-E, 10 min A+F, blue lines), with red light (667 nm, 5 min B-F, 10 min A, red lines), and after dark reversion (1 h D+F, 24 h A+E, 96 h B+C, black lines). The extend of conversion cannot be improved by longer exposure times. The data originate from single sets of measurements (except: *Av*BphP2_D793A dark, black line, Fig. 3E). Manually calculated difference spectra (far-red irradiated minus dark incubated form; dashed lines) are integrated in the graphs. BV IXα was added in excess after cell lysis before purification (except: *Rt*BphP2, coexpression with heme oxygenase *hmuO*, Fig. 3F). The exact Pr/Pfr fractions for each spectrum are given in Tab. 1. The errors for α amount to ±1% for Pfr and ±8% for Pr.

**Table 2:** Compilation of the maximum absorbance wavelength, λ_max_, of the respective pure Pr- and Pfr-forms of the BphPs, of the ratio of the extinction coefficients η = ε(λ_max,Pfr_)/ε(λ_max,Pr_) and of the approximate half-life, t_1/2_, of dark reversion (DR).

| Phytochrome | Pfr $\lambda_{\max}$ | Pr $\lambda_{\max}$ | $\eta$ | DR $t_{1/2}$ |
| --- | --- | --- | --- | --- |
| **PaBphP** | 410 / 752 nm | 398 / 702 nm | 0.65 | ~ 15 min |
| **AtBphP2** | 407 / 753 nm | 394 / 702 nm | 1.1 | < 5 s |
| **AtBphP2_D783N** | 413 / 756 nm | 394 / 701 nm | 1.0 | ~ 5 s |
| **AvBphP2** | 395 / 748 nm | 388 / 698 nm | n.d. | ~ 5 min |
| **AvBphP2_D793A** | 388 / 751 nm | 387 / 698 nm | n.d. | - |
| **RtBphP2** | 405 / 757 nm | 390 / 684 nm | 2.75 | ~ 1 min |
| **XccBphP** | 407 / 754 nm | 392 / 686 nm | 0.76 | 3-4 h |
| **XccBphPΔPAS9** | 414 / 754 nm | 392 / 686 nm | 0.75 | ~ 50 min |

n.d.,: not determined

### Autokinase activity in bathy phytochrome is inhibited in darkness

All bathy phytochromes investigated in this study share the formation of a pronounced Pr- form under far-red light and the accumulation of a Pfr-form in darkness. In order to investigate, whether the observed correlation between absorbing form and autokinase activity in *Pa*BphP is conserved among bathy phytochromes, we investigated the autokinase activity of *At*BphP2_D783N, *Av*BphP2_D793A and *Rt*BphP2 (Fig. 4). They all consistently show reduced autokinase activity in darkness (Fig. 4A-C). While they display similar autophosphorylation signals under far-red, blue, and red-light conditions, they have different calculated Pr/Pfr-fractions. Thus, a strict correlation between autophosphorylation signal under illumination and Pr/Pfr, is not as obvious as for *Pa*BphP. However, we propose that the proportion of active kinase in a given phytochrome can be derived from the spectrally determined Pr/Pfr-fraction, as only the Pr-form is autokinase active.

**Fig. 4:**
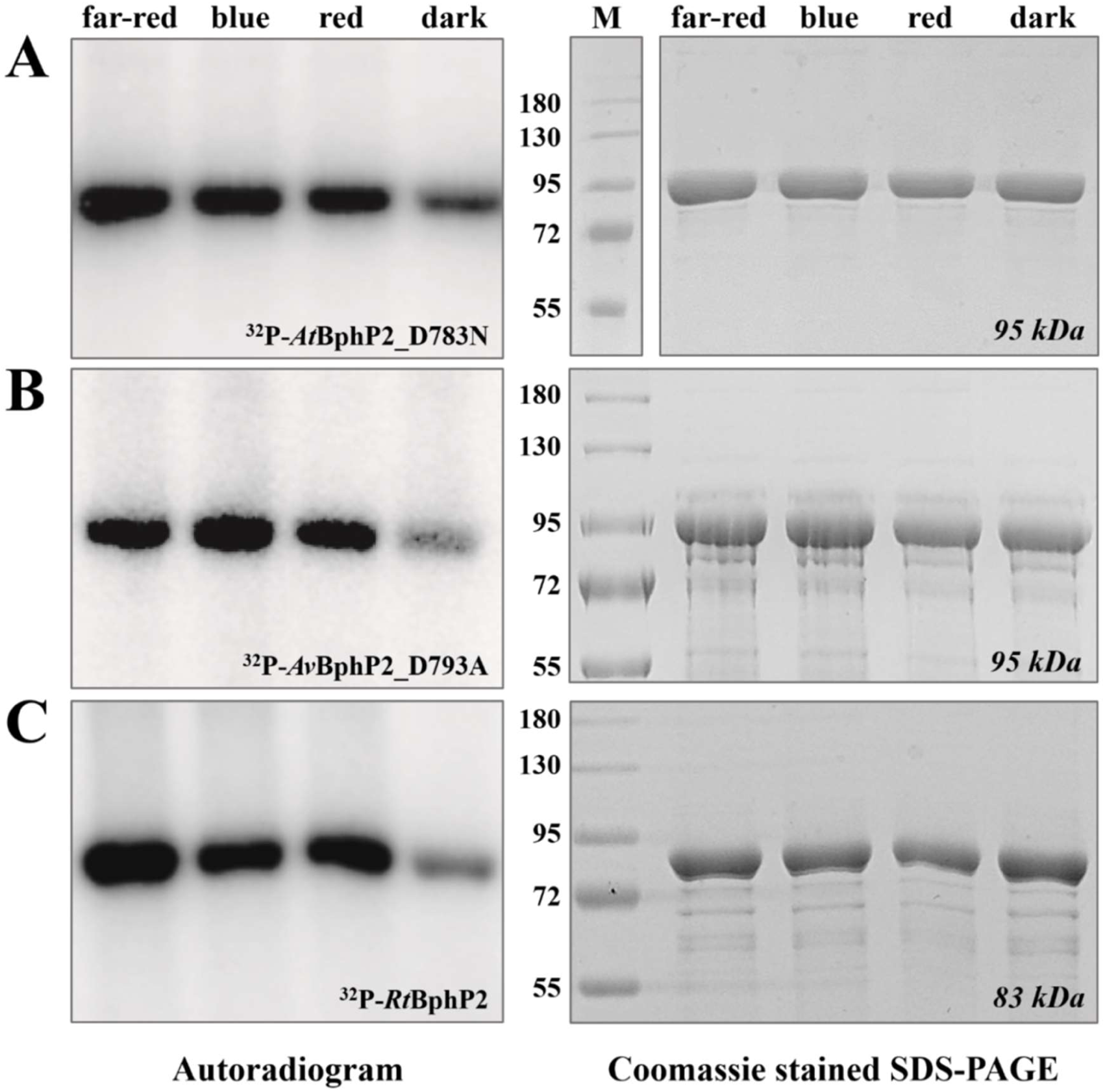
Autophosphorylation activity of recombinant produced *At*BphP2_D783N, *Av*BphP2_D793A and *Rt*BphP2 under different light conditions. Autoradiogram (left) and corresponding Coomassie-stained SDS-PAGE (10 % (w/v), right) gel after *in vitro* radiolabeling of *At*BphP2_D783N **(A)**, *Av*BphP2_D793A **(B)** and *Rt*BphP2 **(C)** with [γ-^32^P]-ATP under far-red (791 nm), blue (426 nm), and red (667 nm) light as well as dark conditions. Autophosphorylation reaction was terminated after 5 min and samples were separated by gel electrophoresis. Proteins were pre-irradiated with the respective light for 5 min or incubated dark overnight before adding the radiolabeled ATP. *At*BphP2_D783N and *Av*BphP_D793A has a relative molecular weight of approximately 95 kDa and *Rt*BphP2 of 83 kDa (M = marker; Color Prestained Protein Standard, Broad Range, NEB).

### Dark reversion

In addition to light-induced photoconversion, phytochromes also undergo a light-independent thermal conversion (a.k.a. dark reversion, DR) (45). This process does influence the amount of available physiologically active phytochromes in the bacterial cell. In bathy phytochromes, DR leads to the accumulation of the kinase inactive Pfr-form (see also Fig. 2). To compare the dark reversion rates, we first irradiated all phytochromes with far-red light to generate pure Pr-forms. The following dark adaptation was tracked by absorption spectroscopy for 24 h (*Pa*BphP, *At*BphP2, *At*BphP2_D783N, *Av*BphP2 as well as *Rt*BphP2) or for 96 h (*Xcc*BphP and *Xcc*BphPΔPAS9), and Pr/Pfr-fractions were calculated for selected timepoints (Fig. 5). The fastest DR was observed for *At*BphP2_D783N (green line) with a half-life of less than 5 sec, which agrees with findings of Piatkevich *et al.*for wildtype *At*BphP2 (46). The kinetics of the respective WT variant, *At*BphP2_D783N (turquoise line), does not vary much, which proves that the exchanged amino acid residue has no effect on the DR process (Tab. 2). The Pr state of *Rt*BphP2 (orange line) is also very unstable and undergoes a fast adaptation in the dark with a half-life of around 1 min. The relaxation progress for *Pa*BphP (black line) is decreased in comparison to the other phytochromes and tenfold slower than the reversion of *Rt*BphP2. The DR rate of *Xcc*BphP (pink line) and *Xcc*BphPΔPAS9 (grey line) is very slow, albeit the formation of the Pfr-state is accelerated for the truncated version. In the study by Otero *et al.,* it was shown that the output module (PAS9 domain) modulates the chromophore regarding the dark adaptation kinetics (32). *Xcc*BphPΔPAS9 has a half-life of around 50 min for the ground state formation whereas the dark reversion of *Xcc*BphP is decelerated (t_1/2_ = 3-4 h) and reaches the maximum Pfr content (72 %) after ∼96 h dark incubation.

**Fig. 5:**
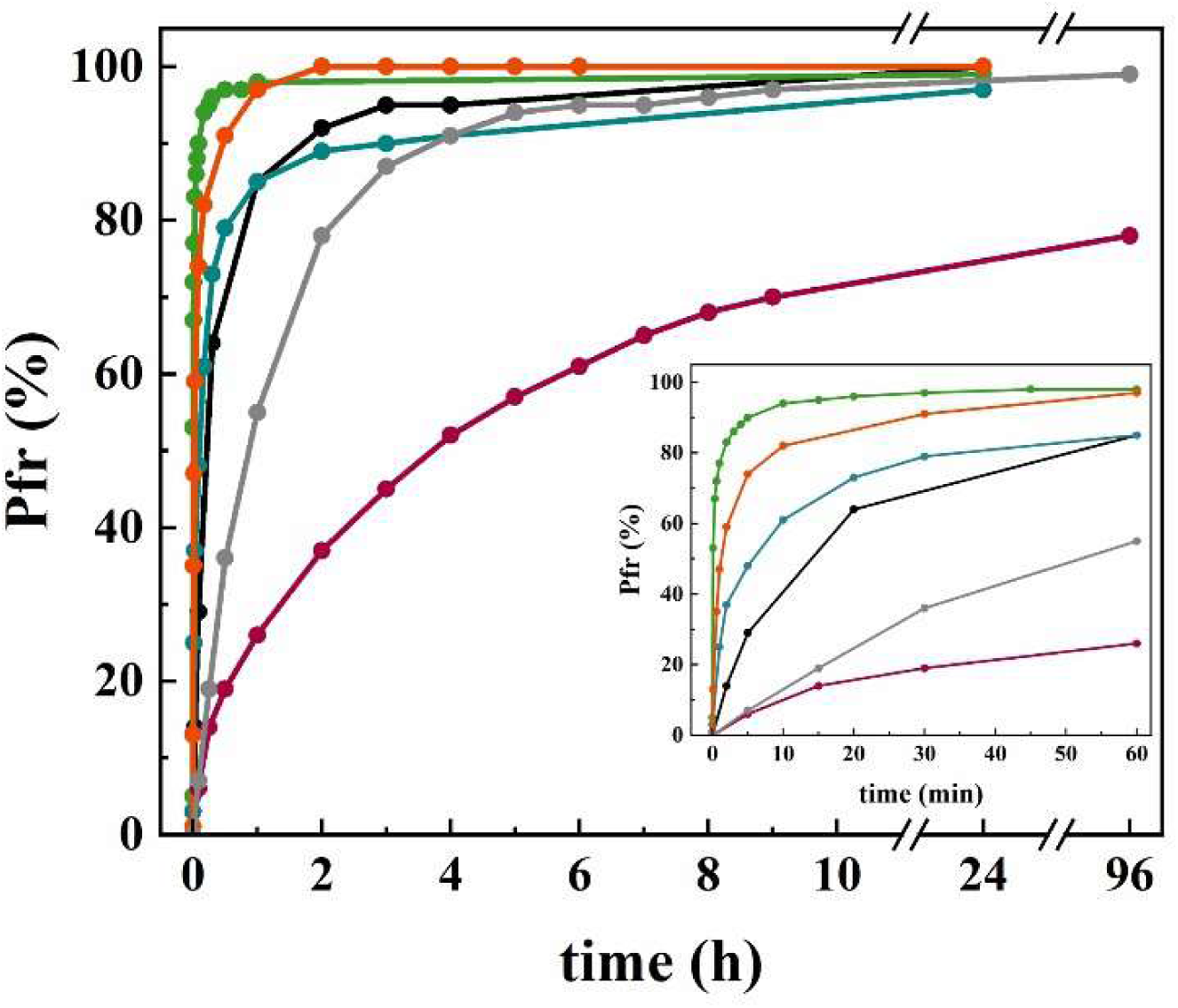
Kinetics of dark reversion of *Pa*BphP, *At*BphP2_D783N, *Av*BphP2, *Rt*BphP2, *Xcc*BphP, and *Xcc*BphPΔPAS9. Sufficient far-red irradiation to allow formation of Pr-form preceded the incubation in darkness. Pr-to-Pfr conversion was monitored for a period of 24 h (*Pa*BphP, black line; *At*BphP2_D783N, green line; *Av*BphP2, turquoise line; *Rt*BphP2, orange line) or 96 h (*Xcc*BphP, pink line; *Xcc*BphPΔPAS9, gray line) and the kinetics were plotted in a line chart. The inset shows the dark reversion up to 60 min. The calculated half-lifes (50 % phytochromes in Pfr-form) are shown in Tab. 2.

## Discussion

### Light activation of bathy phytochromes results in structural rearrangements that lead to autokinase activation

Bacterial bathy phytochromes are a subgroup of phytochrome photoreceptors with a Pfr ground state. By reanalyzing the model bathy phytochrome of *P. aeruginosa*, we were able to establish that a pure Pfr-form is only formed in darkness, while different qualities of light lead to various Pr/Pfr-fractions. Most importantly, using a combination of experimental data and mathematical calculations, we were able to establish a correlation between the Pr/Pfr-fractions under photostationary conditions and the corresponding autokinase activity output. Only the Pr-form contributes to autokinase activity while the Pfr-form represents the inactive state of the kinase. Comparing these results to prototypical bacterial phytochromes with a Pr ground state highlights the similarity between both phytochromes types, where the kinase active form is always the Pr-form (28,29,40–42,47) and that kinase activity is inhibited upon increasing Pfr-proportion (Fig. 6).

**Fig. 6:**
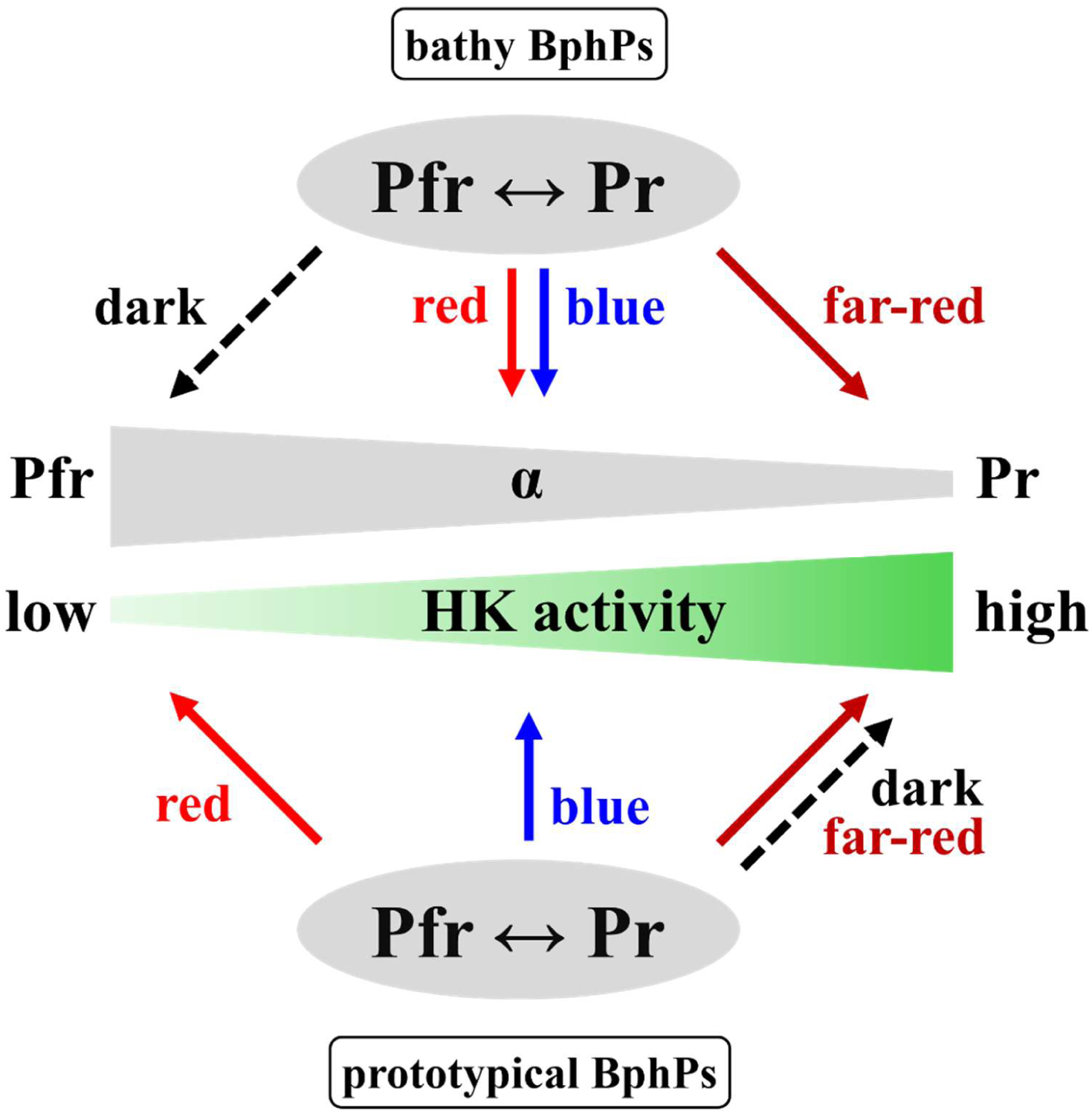
Bathy and prototypical phytochromes act as molecular light switches to adjust their histidine kinase (HK) activity. Bathy and prototypical phytochromes function as switches that regulate the autokinase output in the bacterial cells. It is not a simple on/off system, but rather adjusts HK activity as Pr-fractions increase (= α decreases) and Pfr-fractions decrease (= α increases). Far-red light (for bathy BphPs) and darkness (for prototypical BphPs) are associated with the highest Pr-fractions, lowest α, and highest HK activity. Darkness (for bathy BphPs) and red-light conditions (for prototypical BphPs) are associated with the highest Pfr-fractions, highest α, and lowest HK activity. Blue and red light induce mixed forms with different Pr/Pfr-fractions and kinase activity based on the Pfr-fraction. The α value defines the fraction of Pfr available in a phytochrome population (α = [Pfr]/([Pfr]+[Pr])). The model for the prototypical BphPs is only preliminary because comprehensive investigations on their spectral and autokinase behavior were not conducted in this study.

During the conversion of Pr to Pfr, a significant structural change occurs in the PCM which will immediately impact the HKD (34). A classical bacterial HKD is composed of the dimerization histidine phosphotransfer domain (DHp) and the catalytic adenosine triphosphate–binding domain (CA). The DHp domain is a dimeric four-helix bundle, formed by two helical hairpin monomers (48). The N-terminal DHp helix α1 carries the characteristic H-box sequence motif, where the phospho-accepting His residue is located. The monomeric CA domain adopts an α/β sandwich fold related to known nucleotide-binding folds (48). CA domain sequences exhibit conserved motifs, with residues forming an ATP-binding cavity and a flexible ATP lid. Upon ATP binding and hydrolysis, autophosphorylation in bacterial HKD can occur in *cis* (within one subunit) or *trans* (intersubunit autophosphorylation) via a common mechanism that is independent of directionality of phosphorylation (49–51). Based on recent molecular modeling of full-length *Pa*BphP with the *cis*-directional HKD 853 of *Thermotoga maritima,* it has been proposed that light activation of *Pa*BphP induces a large quaternary structural rearrangement at the dimeric interface resulting in the formation of the Pr-state. This structural change causes a shift in the extended central helical bundles leading to a modification in the quaternary assemblies of the histidine kinase output domain. Upon light-induced structural rearrangement, the helical spines of the DHp domains are twisted together, reducing the spatial gap between them and, concomitantly, putting the CA domains into closer proximity, enabling efficient phosphoryl group transfer as compared to the Pfr state (34). As BphPs exist as dimers, it has been acknowledged in the field that both, Pr/Pr and Pfr/Pfr homodimers, as well as Pr/Pfr heterodimers are feasible (18,35,36,38). It is currently unclear whether Pr/Pr homodimers are the only kinase active form, or if there are Pr/Pfr heterodimers with “half site reactivity”. In our opinion, the latter is likely only possible for the *cis*-directionality of phosphorylation. Consequently, future combination of improved structural analysis and biochemical studies will be necessary to resolve this intriguing question.

### The protein conformation adopted in darkness is the switch-off signal for bathy BphPs

One significant difference between prototypical and bathy BphPs is the input signal required to turn the kinase off (i.e., to develop the Pfr-form) which is red light for the prototypical phytochromes and darkness for the bathy ones (Fig. 6). DR in the case of bathy BphPs function as a mechanism for the organism to switch off the system. The time frame for the formation of a pure Pfr-form subsequent to transfer into darkness can fluctuate greatly, ranging from mere seconds to several hours, or even days (15,26). The DR rate can also be affected by various cellular conditions such as temperature or pH (15,52–54). For *At*BphP2, it was demonstrated that the same proton translocations that are responsible for the formation of the photoactivated state and activating the output module, are also involved in the thermal back-isomerization of the chromophore. Thus, the identical re-arrangement of protons also stimulates the deactivation of the output/transmitter module, similar to a negative feedback mechanism (54). UV/Vis spectroscopy was utilized to determine the approximate half-life of DR, indicating the time point where 50 % of the bathy BphP population is present in its Pfr-form. Compared to the 15 min half-life DR of *Pa*BphP, the other examined bathy BphPs exhibit notably shorter DR times. However, *Xcc*BphP is an outlier, requiring 1-4 h to achieve 50 % of its Pfr-form. Currently, it is completely unclear whether there is a physiological reason behind the time required to return to the ground state. It can only be speculated whether the combination of DR and temperature sensing has a biological impact, or whether a rapid DR compensates for phosphatase activity. Bacterial histidine kinases often possess intrinsic phosphatase activity, or an external phosphatase is included in the signal transduction system (like KinB in the *Pa*BphP-AlgB system (43)). Phosphatases enable the organism to accelerate the rate of response regulator dephosphorylation to reset the system and to prevent crosstalk (55). Bathy phytochrome two-component systems lacking a phosphatase may rely on rapid DR to efficiently return to their inactive and unphosphorylated state. While *Pa*KinB is described as a phosphatase in the regulatory system with *Pa*BphP and *Pa*AlgB (43), there is no literature supporting the existence of phosphatases in the other bathy phytochrome systems.

### Bathy BphPs are involved in regulating virulence

Based on the results presented in this work, it can be concluded that bathy phytochromes function as light/dark sensors rather than classical far-red/red light sensors. Changing the Pr/Pfr-fractions enables the organisms to adjust their autokinase output in response to changing light qualities and environments. The system can only be turned off by darkness and the related pure or highly enriched Pfr-form. Organisms that possess only a bathy BphP as their single phytochrome photoreceptor are capable of utilizing various light qualities to fine-tune their physiological output response by modulating the proportion of active Pr kinase. For organisms such as *A*. *tumefacience*, *A. vitis* and *R. tataouinensis* that possess a prototypical and a bathy phytochrome, both BphPs have opposing activities. This enables the bacteria to further enhance their adaptation to environments abundant in red and far-red light (29,31,33).

Although a physiological response to light has been reported for all organisms containing bathy phytochromes, the detailed molecular mechanism is often still not fully understood (33,56–58). For *Pa*BphP it has been demonstrated previously that upon autophosphorylation (i.e. high Pr-fraction), the phosphoryl group is transferred to the corresponding response regulator *Pa*AlgB. In turn, phosphorylated *Pa*AlgB (indirectly) inhibits swimming motility, biofilm formation, and the secretion of the virulence factor pyocyanin (43). The whole system is furthermore controlled by the phosphatase *Pa*KinB, which is responsible for the dephosphorylation of *Pa*AlgB and thus acts as an antagonist to *Pa*BphP. Therefore, it appears that the Pfr-form of the BphPs linked to darkness induces the derepression of virulence gene expression in both plant and human pathogens (43,56,57). Essentially, it is reasonable for *P. aeruginosa* not to inhibit biofilm formation and production of virulence factors in the absence of light. The lack of light inside the human body can promote infections in the lungs or other human tissues. Additionally, the bacteria have the ability to sense light to detect the highest efficacy of the host immune system, making it advantageous for the pathogen to infect the host at night to evade their defenses (43). Effective infections of the plant pathogens *X. campestris*, *A. tumefaciens*, and *A. vitis* are also facilitated by maintaining virulence and flagella synthesis in the absence of light. The defense mechanism does not require evasion during the night as it is regulated by light and reaches its peak functionality during the day (56). Nevertheless, it remains a fine-tuning system dependent on the incident light rather than a simple on/off system.

Although our data provide some more insights into the signal transduction within bathy phytochromes, further research is required to ultimately understand the molecular mechanisms of these fascinating photoreceptors and their function in bacteria.

## Experimental Procedures

### Bacterial Strains, Media, and Growth Conditions

The *P. aeruginosa* type strain UCBPP-PA14 and *E. coli* BL21(DE3) were from our laboratory stock and were used as the initial strains for the main part of the experiments. All further strains used in this study were stored in glycerol (*E. coli*) or DMSO (*P. aeruginosa*) stocks and are listed together with the plasmids in Table S2. *E. coli* strains, *P. aeruginosa* strains, and mutants were cultured in lysogeny broth (LB-Lennox; 10 g/l tryptone, 5 g/l NaCl, 5 g/l yeast extract) or on LB agar (15 g/l) at 37 °C. When required, antibiotics were used at the following concentrations: 100 µg/ml ampicillin (*E. coli*), 50 µg/ml kanamycin (*E. coli*), 300 µg/ml gentamycin (*P. aeruginosa*) and 100 µg/ml tetracycline (*P. aeruginosa*).

### Construction of Mutant Strains and Plasmids

In course of this work the two-step allelic exchange method was used for *P. aeruginosa* genome engineering (59). This method was applied to generate markerless in-frame chromosomal deletions in PA14. To construct gene knockouts, DNA fragments flanking the gene of interest were amplified from genomic DNA of PA14 and ligated into the suicide vector pEXG2 (pEXG2_Δ*bphP*, pEXG2_Δ*bphOP*). The created plasmids were used to transform *E. coli* S17-I and mobilized by conjugation into the recipient strain PA14 via biparental mating. Single-crossover recombinants with the site-specifically integrated plasmid, resulting from homologous recombination, were selected on LB agar containing gentamycin. Incubation of the merodiploid exconjugants on LB agar overnight forces the removal of the vector DNA containing the *sacB* gene. Double-crossover mutants were recovered using counter-selection on LB agar containing 15 % sucrose. Mutants were confirmed by PCR as well as by Sanger sequencing (60,61).

The *P. aeruginosa bphP* gene (NCBI reference sequence: NP_252806.1) was PCR-amplified from PAO1 genomic DNA, where it has a 99 % similarity to PA14. The fragment was ligated into the pHERD26T (62) expression vector via restriction sites *Xba*I/*Kpn*I to obtain pHERD_*Pa*BphP. QuikChange site-directed mutagenesis of *PabphP* (H513A, D194H, S261A) was performed using pHERD_*Pa*BphP. The resulting ORFs encode *Pa*BphP or the *Pa*BphP derivatives under the control of an *ara* promoter with a C-terminal Strep-tag II.

The plasmid pSA2 (= pET21b_*At*BphP2) (63) and pETavi3496 (= pET21b_*Av*BphP2) (31) were kindly provided by T. Lamparter, KIT, Germany. QuikChange site-directed mutagenesis of *AtbphP*2 and *AvbphP*2 was performed using these two plasmids to obtain pET21b_*At*BphP2_D783N and pET21b_*Av*BphP2_D793A, respectively. The plasmid pBAD/HisB-*Rt*BphP2-HmuO was kindly provided by G. De Luca, Aix-Marseille Université, France (33) and pET24a_*Xcc*BphP as well pET24a_*Xcc*BphPΔPAS9 (32) were kindly provided by H. Bonomi, Fundación Instituto Leloir, Argentina. All encoded proteins were fused with a 6x polyhistidine-tag and the gene expression is under control of a T7 (pET21b, pET24a) or an *ara* (pBAD/HisB) promoter.

All sequences of oligonucleotides used, are listed in Table S3.

### Protein Production and Purification

*P. aeruginosa* PA14Δ*bphP* cells harboring pHERD_*Pa*BphP were grown in 50 ml LB media supplemented with tetracycline overnight at 37 °C. For protein production the cells were used to inoculate 1 l LB-tetracycline media and gene expression was induced by addition of 0.1 % L(+)-arabinose at 0.7 OD_600_. For producing the apo-phytochrome pHERD_*Pa*BphP was mobilized by conjugation into PA14Δ*bphOP* and followed the same procedure as before. *E. coli* BL21(DE3) cells harboring pET21b_*At*BphP2, pET21b_*At*BphP2_D783N, pET21b_*Av*BphP2, pET21b_*Av*BphP2_D793A, pET24a_*Xcc*BphP or pET24a_*Xcc*BphPΔPAS9 and *E. coli* Top10 cells harboring pBAD/HisB-*Rt*BphP2-HmuO were grown in 15 ml LB media supplemented with ampicillin (pET21b, pBAD/HisB) or kanamycin (pET24a) overnight at 37 °C. For protein production the cells were used to inoculate 1 l LB-ampicillin or LB-kanamycin, respectively. The expression of the phytochrome genes - for pBAD/HisB also for the heme oxygenase gene - were induced by addition of 0.5 mM isopropyl β-D-1-thiogalactopyranoside (IPTG) (pET21b, pET24a) or 0,1 % L(+)arabinose (pBAD/HisB) at 0.5 OD_600_. After incubation for further 5 d at 10 °C (*Av*BphP2 and the variant), 22 h at 17 °C (*Pa*BphP, *At*BphP2, *Xcc*BphP and the respective variants) or 4 h at 30 °C (*Rt*BphP2) the cells were harvested by centrifugation for 15 min and dissolved in lysis buffer (100 mM Tris-HCl [pH 8.0], 300 mM NaCl; for *Rt*BphP2, *Xcc*BphP and *Xcc*BphPΔPAS9 1 % Triton-X-100; 2 ml buffer per g of cells) supplemented with lysozyme and DNase I. The cells were lysed by cell fluidizer (Microfluidics™) technology. Lysed cells were centrifuged and the protein solutions with *Pa*BphP, *At*BphP2, *Av*BphP2, *Xcc*BphP or the protein variants in the supernatant were incubated with excess BV (200 mM) at 4 °C in the dark for an hour to obtain almost complete saturation with the chromophore. The resulting protein solutions with holophytochrome complexes of *Pa*BphP and variants were subjected to Strep-Tactin® Sepharose® resin and treated according to the instructions of the manufacturer. The proteins were eluted with 25 mM d-Desthiobiotin. The protein solutions with holophytochrome complexes of *At*BphP2, *Av*BphP2, *Xcc*BphP or the variants and the *Rt*BphP2 extract without addition of BV were purified using TALON®Superflow™ (Cytiva) according to the instructions of the manufacturer and were eluted with 75 mM imidazole. After elution, the purified proteins were dialyzed in kinase buffer (50 mM Tris-HCl [pH 8.0], 100 mM NaCl, 5 mM MgCl_2_, 10 % (v/v) glycerol) overnight and concentrated using a centrifugal concentrator (Amicon). To assess purity, the proteins were analyzed by Coomassie brilliant blue stained SDS-PAGE gels. Protein concentration was estimated using the calculated molar extinction coefficient at 280 nm provided by Protein Calculator v3.4 (http://protcalc.sourceforge.net/) based on the amino acid sequence with the respective tag.

### UV/Visible Spectroscopy

UV/Vis measurements were performed on an Agilent 8453 UV/Visible spectroscopy system at room temperature in a Hellma precision quartz cuvette (100 µl sample volume, 10 mm optical path). For spectroscopy the purified proteins were diluted in 1x kinase buffer.

Photostationary states of phytochrome were obtained by exposing the sample in the spectrometer chamber sufficiently long to darkness, far-red (791 ±2 nm, 200 mW, 61 mW/cm^2^, *Panasonic* LNCT28PS01WW, laser diode), red (667 ± 2 nm, 100 mW, 27 mW/cm^2^, *Panasonic* LNCT28PS01WW, laser diode) or blue (426 ± 70 nm, 0.65 mW, 0.4 mW/cm^2^, *Everlight* DLE-038-046, LED) light irradiation (emission spectra cf. Fig. S4). It was ensured that the variations in the listed intensities (e.g. by slightly different illumination geometries) that occurred in the course of the work on the spectrometer and the kinase assay (see below) had no effect on the photostationary Pr/Pfr equilibrium determined.

To measure the dark reversion rates, we first irradiated all phytochromes with far-red light to generate pure Pr-forms. The following dark adaptation was tracked by absorption spectroscopy for 24 h (*Pa*BphP, *At*BphP2, *At*BphP2_D783N, *Av*BphP2, and *Rt*BphP2) or for 96 h (*Xcc*BphP and *Xcc*BphPΔPAS9).

### Calculation of Pr and Pfr fraction in (photo-)stationary phytochrome samples from absorption spectra

To determine the fraction α = [*P_fr_*]/([*P_fr_*] + [*P_r_*]) of Pfr in a given (photo-)stationary phytochrome sample, the Q-band region in the respective UV/Vis spectrum was analyzed. To this end, the characteristic photocycle of the respective phytochrome was reduced to a simple binary kinetic model, comprising the two-limiting species Pfr and Pr, two photochemical rates k_p_fr_→P_r_ and_ k_p_r_→P_fr__, and the rate for dark-reversion from Pr to Pfr, k_DR_. Based on this model, α for any given (photo-)stationary phytochrome sample can be obtained by simulating (Matlab 2021b) the experimental Q-band spectrum A(ṽ) via the fitting function 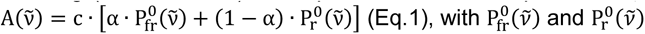 the pure spectra of Pfr and Pr, respectively, taken at the same concentration to achieve the quantitative relation between their extinction coefficients, and c an overall scaling constant. P_fr_^0^(*ṽ*) was obtained by complete dark-reversion. P_r_^0^(*ṽ*) was obtained by illuminating the sample in the far-red before and during the absorption measurement (inhibiting dark reversion). To account for scattering contributions, a linear background was subtracted from the Q-band region (ca. 550 - 850 nm). For the fitting procedure, P_fr_^0^(*ṽ*) and P_r_^0^(*ṽ*) were modelled as superpositions of multiple Gaussian functions. For the stationary spectra and related fits cf. Fig. S6, Table S1. The errors for α amount to ±1% for Pfr and ±8% for Pr.

### *In vitro* kinase assay of phytochrome proteins with well-defined Pfr fraction

Autophosphorylation assays were performed in kinase buffer with purified full-length *Pa*BphP, *At*BphP2_D783N, *Av*BphP2_D793A, and *Rt*BphP2 in a newly designed illumination setup (Fig. S5). Autophosphorylation reaction was initiated by addition of 2.5 µCi [γ-^32^P]-ATP (Hartmann Analytic) to 5 µM purified phytochrome in a final volume of 12.5 µl. *Pa*BphP, *At*BphP2_D783N, *Av*BphP2_D793A, and *Rt*BphP2 were irradiated 5 min with far-red, red, blue light or incubated dark overnight before addition of radiolabeled ATP. The start of the reaction with the ATP was followed by another 5 min incubation under the respective light condition or in darkness before termination. To stop the reactions 2.5 µl 4x SDS sample buffer and 1.5 µl 0.5 M EDTA were added. Samples were separated on a 10 % SDS-PAGE gel and imaged using a Typhoon™ FLA 7000 (GE Healthcare). Irradiation intensity as well as time, and corresponding definition of α-values were determined by previous UV/Vis spectroscopy. Light exposure for kinase assays used the same laser diodes and LED as well comparable illumination geometry as utilized for the spectroscopy experiments.

## Data availability

All data are contained within this manuscript. All described strains and plasmid constructs are available upon request from the corresponding author.

## Supporting information

This article contains supporting information.

Tables S1-S3

Figures S1-S6

## Acknowledgments

We are very grateful to T. Lamparter (Karlsruhe Institute für Technologie, Germany), G. De Luca (Aix-Marseille Université, France) and H. Bonomi (Fundación Instituto Leloir, Argentina) for gift of plasmids used in this study. Furthermore, we thank J. Clark Lagarias for helpful discussion.

## Author Contributions

NFD, RD, JP and CH designed the study, CH, MS, IS, JP, XM performed the research, CH, MS, JP, JH, RD and NFD evaluated data, JH contributed technological expertise, CH and NFD wrote manuscript draft with help of all authors.

## Funding and additional information

This work was in part funded by the Landesforschungsschwerpunkt Rheinland-Pfalz BioComp.

## Conflict of interest

There are no competing interests.

## Notes

### Competing Interest Statement

The authors have declared no competing interest.

