## supplemental material for "Darkness inhibits autokinase activity of bacterial bathy phytochromes"

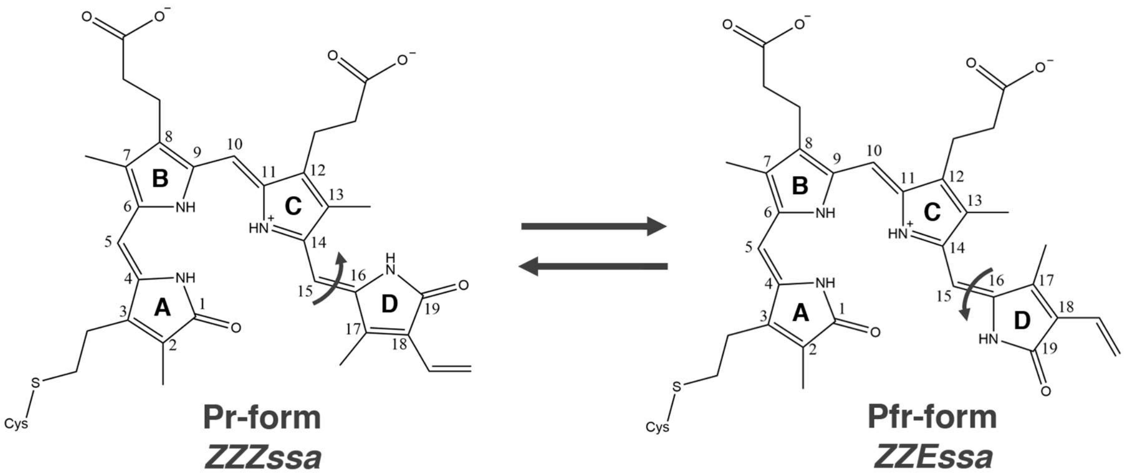

**Fig. S1.** Conformational structure of the chromophore Biliverdin IX$\alpha$ in the Pr-form (*ZZZssa*) and Pfr-form (*ZZEssa*).

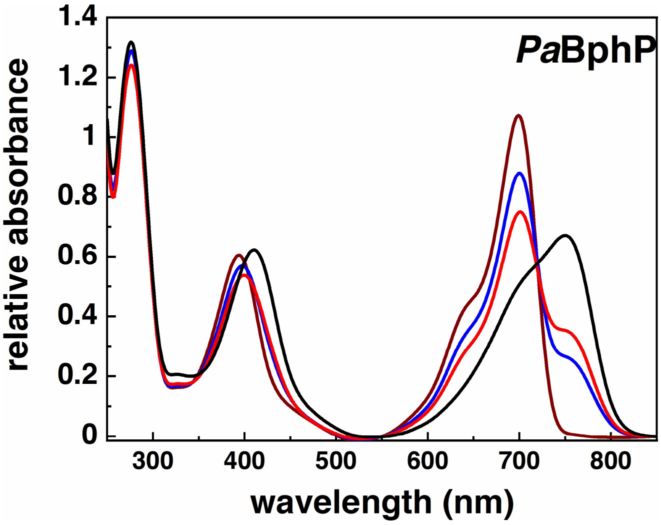

**Fig. S2.** Absorbance spectra of *Pa*BphP (smoothed with OriginLab, Version 2022, OriginLab Corporation, Northampton MA, USA. “Loess Use Proportion for Span, Span 0.1) after illumination with far-red light (791 nm, 5 min, far-red line), with blue light (426 nm, 10 min, blue line), with red light (667 nm, 10 min, red line) and after dark reversion (24 h, black line).

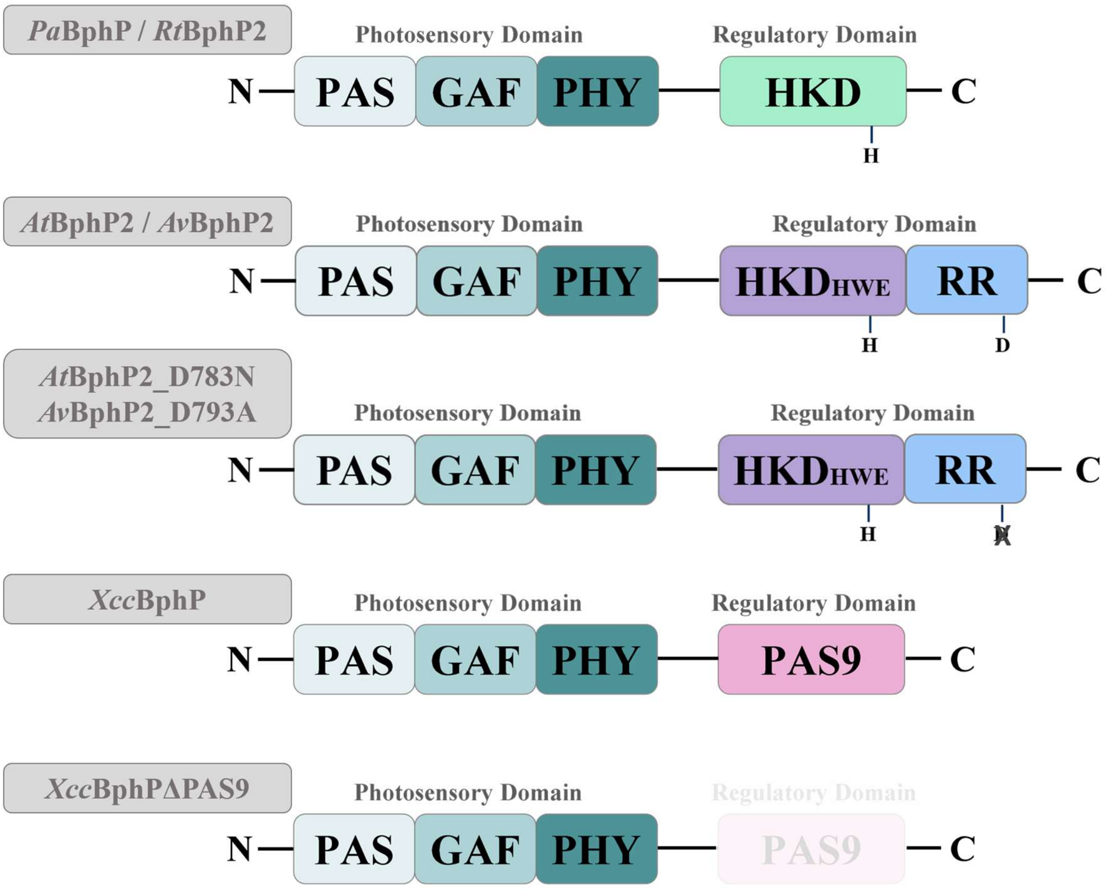

**Fig. S3.** Domain organization of bathy phytochromes investigated in this study.

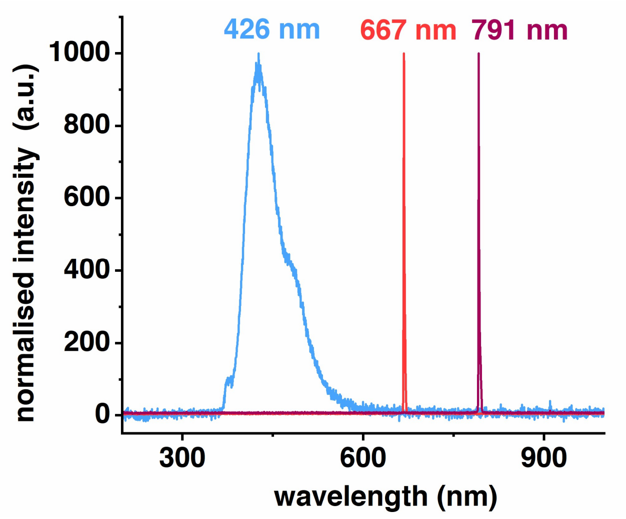

**Fig. S4.** Normalized emission spectra of light sources used for illumination of the samples, obtained with an optical multichannel analyzer (OMA)(*Ocean Optics* OMA, USB2000+UV/Vis).

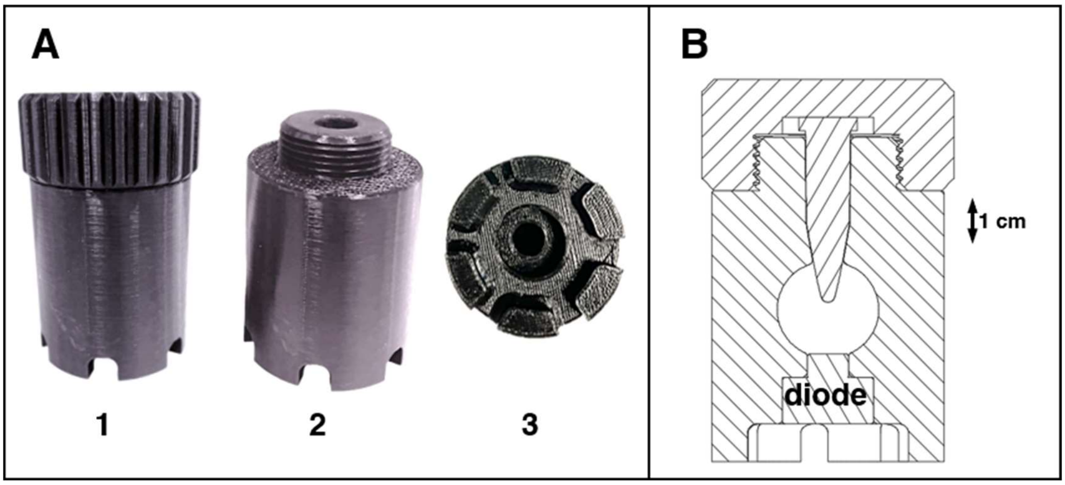

**Fig. S5. Photograph (A) and technical drawing (B) of the new illumination set-up used for the radioactive kinase assays.** It consists of a main pod (A2) with a screw-cap (A1), both fashioned from black polylactide, and thus protects the sample from unwanted ambient light. A diode can be inserted from the bottom (A3, B). The phytochrome sample itself is located in an Eppendorf tube (B).

**Simulation of the pure form spectra by sum of Gaussians.**

The experimental pure form spectra ($P_{fr}^0(\tilde{\nu})$ and $P_r^0(\tilde{\nu})$) of the Pfr and Pr form of the various phytochromes, were simulated (after background subtraction) via $P^0(\tilde{\nu}) = \sum_i G_i(\tilde{\nu})$ (Eq. S1), with $G_i(\tilde{\nu}) = y_i^0 + a_i/(w_i\sqrt{\pi/2}) \cdot \exp(-2 \cdot (\tilde{\nu} - \tilde{\nu}_i^c)^2/w_i^2)$ (Eq. S2), $y_i^0$ set to zero, $a_i$ the area, $w_i$ the width, and $\tilde{\nu}_i^c$ the center wavenumber (1).

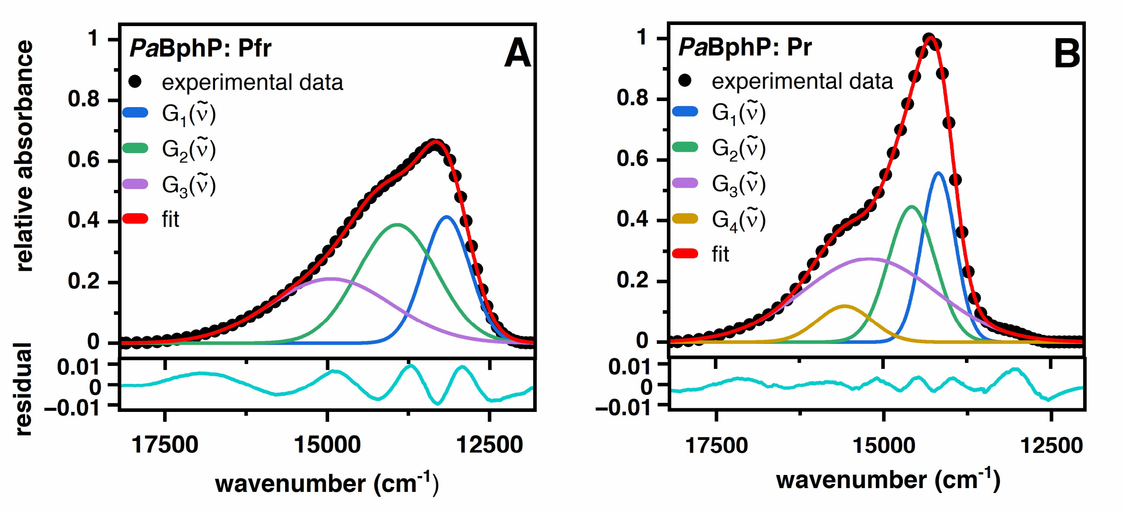

**Fig. S6.** Experimental pure form spectrum $P_{fr}^0(\tilde{\nu})$ (left) and $P_r^0(\tilde{\nu})$ (right) of *Pa*BphP, simulated (OriginLab 2022b) by a sum of three and four Gaussian functions, respectively (cf. Eq. S2). The fit parameters are given in Table S1.

**Table S1.** Fit parameters for the simulation of the pure form spectra $\boldsymbol{P^0_{fr}(\tilde{\nu})}$ and $\boldsymbol{P^0_{r}(\tilde{\nu})}$ of the investigated bathy-phytochromes. 1: *Pa*BphP, 2: *At*BphP2 WT, 3: *At*BphP2_D783N, 4: *Av*BphP2_WT, 5: *Av*BphP2_D793A, 6: *Rt*BphP WT, 7: *Xcc*BphP WT, 8: *Xcc*BphPΔPAS9

| | | $\mathbf{P^0_{fr}(\tilde{\nu})}$ | | | | $\mathbf{P^0_{r}(\tilde{\nu})}$ | | | |
|---|---|---|---|---|---|---|---|---|---|
| | | $G_1$ | $G_2$ | $G_3$ | $G_4$ | $G_1$ | $G_2$ | $G_3$ | $G_4$ |
| **1** | $\tilde{\nu}^c$ | 13165 ± 3 | 13926 ± 16 | 14946 ± 108 | - | 14172 ± 4 | 14572 ± 29 | 15208 ± 4 | 15572 ± 13 |
| | $w$ | 721 ± 9 | 1216 ± 54 | 1879 ± 75 | - | 503 ± 8 | 704 ± 30 | 2001 ± 13 | 812 ± 21 |
| | $a$ | 375 ± 21 | 594 ± 78 | 499 ± 60 | - | 351 ± 35 | 392 ± 40 | 687 ± 12 | 121 ± 8 |
| **2** | $\tilde{\nu}^c$ | 12956 ± 7 | 13196 ± 18 | 13936 ± 20 | 15195 ± 71 | 14059 ± 30 | 14422 ± 85 | 14859 ± 19 | 15771 ± 193 |
| | $w$ | 534 ± 17 | 786 ± 14 | 1389 ± 21 | 1700 ± 52 | 492 ± 23 | 606 ± 65 | 1825 ± 44 | 2690 ± 89 |
| | $a$ | 177 ± 28 | 492 ± 38 | 1215 ± 49 | 529 ± 47 | 234 ± 82 | 254 ± 85 | 1046 ± 88 | 428 ± 90 |
| **3** | $\tilde{\nu}^c$ | 12988 ± 2 | 13297 ± 4 | 14056 ± 22 | 15027 ± 75 | 14164 ± 4 | 17289 ± 34 | 14659 ± 11 | 15037 ± 6 |
| | $w$ | 565 ± 4 | 883 ± 10 | 1411 ± 37 | 2156 ± 42 | 523 ± 5 | 1237 ± 55 | 481 ± 13 | 1968 ± 8 |
| | $a$ | 204 ± 6 | 598 ± 31 | 837 ± 68 | 513 ± 42 | 407 ± 7 | 56 ± 4 | 124 ± 8 | 1229 ± 3 |
| **4, 5** | $\tilde{\nu}^c$ | 13133 ± 2 | 14194 ± 2 | 15155 ± 21 | - | 14139 ± 3 | 14490 ± 20 | 15226 ± 5 | - |
| | $w$ | 756 ± 2 | 1090 ± 7 | 1963 ± 18 | - | 487 ± 10 | 737 ± 16 | 1872 ± 4 | - |
| | $a$ | 421 ± 3 | 660 ± 14 | 741 ± 15 | - | 245 ± 24 | 411 ± 27 | 1098 ± 5 | - |
| **6** | $\tilde{\nu}^c$ | 13009 ± 2 | 13396 ± 14 | 14300 ± 22 | 14677 ± 18 | 14398 ± 2 | 15498 ± 11 | - | - |
| | $w$ | 538 ± 6 | 890 ± 8 | 908 ± 25 | 1936 ± 13 | 971 ± 7 | 2453 ± 12 | - | - |
| | $a$ | 677 ± 34 | 2059 ± 41 | 735 ± 56 | 2021 ± 43 | 744 ± 11 | 1720 ± 14 | - | - |
| **7** | $\tilde{\nu}^c$ | | | | | 14551 ± 4 | 15082 ± 9 | 15783 ± 2 | 15783 ± 11 |
| | $w$ | from ΔPAS9 scaled with 1.01 | | | | 642 ± 3 | 633 ± 12 | 2493 ± 6 | 1023 ± 14 |
| | $a$ | | | | | 592 ± 10 | 232 ± 14 | 774 ± 4 | 290 ± 9 |
| **8** | $\tilde{\nu}^c$ | 13049 ± 1 | 13616 ± 3 | 14049 ± 4 | 15080 ± 36 | 14574 ± 1 | 15130 ± 3 | 15692 ± 8 | 15897 ± 3 |
| | $w$ | 739 ± 2 | 553 ± 4 | 1419 ± 13 | 2161 ± 22 | 660 ± 2 | 539 ± 6 | 1124 ± 12 | 2453 ± 6 |
| | $a$ | 484 ± 4 | 82 ± 2 | 713 ± 21 | 424 ± 16 | 661 ± 3 | 137 ± 6 | 378 ± 8 | 690 ± 4 |

**Table S2.** Strains and plasmids used in this study.

| Strain or plasmid | Sequence or Characteristics | Reference |
|---|---|---|
| *E. coli* strains | | |
| DH5α | $F^-$ *endA1 glnV44 thi-1 recA1 relA1 gyrA96 deoR nupG purB20* φ80d*lacZ*ΔM15 Δ(*lacZYA-argF*)U169, *hsdR17*($r_K^- m_K^+$), $\lambda^-$ | (2) |
| BL21(DE3) | $F^-$ *ompT gal dcm lon hsdS*$_B$($r_B^- m_B^-$) λ(DE3 [*lacI lacUV5-T7p07 ind1 sam7 nin5*]) [*malB*$^+$]$_{K\text{-}12}$($\lambda^S$) | (3) |
| S17-I | $Tp^r$ $Sm^r$ *recA thi pro hsdR*$^-$*M*$^+$ RP4 : 2-Tc : Mu : Km Tn*7* λpir | (4) |
| Top10 | $F^-$ *mcrA* Δ(*mrr-hsdRMS-mcrBC*) φ80*lacZ*Δ*M15* Δ*lacX74 nupG recA1 araD139*Δ(*ara-leu*)7697 *galE15 galK16 rpsL*($Str^R$) *endA1* $\lambda^-$ | Invitrogen |
| *P. aeruginosa* strains | | |
| PAO1 | *P. aeruginosa* wild type DSM-22644 | (5) |
| PA14 | *P. aeruginosa* wild type UCBPP-14 | (6) |
| PA14Δ*bphP* | 2.0-kb in-frame deletion of *bphP* | This study |
| PA14Δ*bphOP* | 2.7-kb in-frame deletion of *bphOP* | This study |
| Plasmids | | |
| pHERD26T | *E. coli* / *P. aeruginosa* shuttle vector, homologous overexpression in *P. aeruginosa*, $P_{BAD}$, $Tet^R$ | (7) |
| pHERD_*Pa*BphP | pHERD26T derivative, coding region of *bphP* from *P. aeruginosa* (PAO1_4117) at *Xba*I/*Kpn*I site with C-terminal Strep-tag II, $P_{BAD}$, $Tet^R$ | This study |
| pEXG2 | Allelic exchange vector for construction of marker-less deletion mutants in *P. aeruginosa*, $Gm^R$ | (8) |
| pEXG2_Δ*bphP* | pEXG2 derivative, truncated version of *bphP* from *P. aeruginosa* (PAO1_4117; 147 bp) with 621 bp upstream and 818 bp downstream at *Hind*III/*EcoR*I site, $Gm^R$ | This study |
| pEXG2_Δ*bphOP* | pEXG2 derivative, truncated version of *bphOP* from *P. aeruginosa* (PA14_10700/10710; 123 bp) with 513 bp upstream and 451 bp downstream at *EcoR*I/*BamH*I site, $Gm^R$ | This study |
| pET21b_*At*BphP2 | pET21b derivative, coding region of *bphP*2 from *A. tumefaciens* at *BamH*I/*Nde*I site with C-terminal His-tag, T7 promoter, $Amp^R$ | (9) |
| pET21b_*At*BphP2_D783N | pET21b_*At*BphP2 with encoded amino acid exchange → Asp (783) by Asn | This study |
| pET21b_*Av*BphP2 | pET21b derivative, coding region of *bphP*2 from *A. vitis* at *Nde*I/*Xho*I site with C-terminal His-tag, T7 promoter, $Amp^R$ | (10) |
| pET21b_*Av*BphP2_D793A | pET21b_*Av*BphP2 with encoded amino acid exchange → Asp (793) by Ala | This study |
| pBAD/HisB_*Rt*BphP2 HmuO | pBAD/HisB derivative, coding region of *bphP*2 from *R. tataouinensis* (Rta_28950) at *Bgl*II/*EcoR*I site with N-terminal His-tag and *hmuO* from *Bradyrhizobium* sp. ORS278 at *EcoR*I/*Hind*III site, $P_{BAD}$, $Amp^R$ | (11) |
| pET24a_*Xcc*BphP | pET24a derivative, coding region of *bphP* from *X. campestris* pv. *campestris* strain 8004 residue | (12) |

| | | |
|---|---|---|
| | 1-634 (XC_4241) at *Nde*I/*BamH*I site with N-terminal His-tag, T7 promoter, Kan$^R$ | |
| pET24a_*Xcc*BphPΔPAS9 | pET24a derivative, coding region of *bphP* from *X. campestris* pv. *campestris* strain 8004 residue 1-511 (XC_4241) at *Nde*I/*BamH*I site with N-terminal His-tag, T7 promoter, Kan$^R$ | (12) |

**Table S3.** Oligonucleotides used in this study.

| Primer | Sequence |
|---|---|
| pEXG2Δ*bphP*_upF | TTAGCTAAGCTTATGTCCCCATCTCCATCGCCA |
| pEXG2Δ*bphP*_upR | GTACAGGCCCGGGGTGATGCTCGTCAT |
| pEXG2Δ*bphP*_downF | ATCACCCCGGGCCTGTACATCTCCCAG |
| pEXG2Δ*bphP*_downR | AATCTAGAATTCGAACGGCTGGCGTACTTC |
| pEXG2Δ*bphOP*_upF | GCGAATTCGGGCCTGAAGGAAGTGAAGCAGT |
| pEXG2Δ*bphOP*_upR | CGCAGGCAGAAGGTGATTTCTGCGTGCAGGTCACGGG |
| pEXG2Δ*bphOP*_downF | ATCACCTTCTGCCTGCGCCT |
| pEXG2Δ*bphOP*_downR | GCGGATCCGCGTACGGTCTTGCGCTTGC |
| pEXG2_seqF | CGACCTCATTCTATTAGACTCTCGTTTGGATTGC |
| pEXG2_seqR | GTTCGCTCGCGTATCGGTGATTCATTCTG |
| Δ*bphOP*_seqF | GGGGATATCCATTTCGCGGAAG |
| Δ*bphOP*_seqR | GCTCCATGAATTCGTCGGGG |
| pHERD-*Pa*BphP_fwd | CGTCTAGACATGACGAGCATCACCCCGGTTACC |
| pHERD-*Pa*BphP _rev | CCGGTACCGTTCAGGACGAGGAGCCGGTCTCC |
| pHERD-*Pa*BphPH513A_fwd | GCGGTGCTCGGCGCCGACCTGCGCAAC |
| pHERD-*Pa*BphPH513A_rev | GTTGCGCAGGTCGGCGCCGAGCACCGC |
| pHERD-*Pa*BphPD194H_fwd | GCAACGCTACCCGGCCTCGCACATCCCGGCCCAGGCG |
| pHERD-*Pa*BphPD194H_rev | CGCCTGGGCCGGGATGTGCGAGGCCGGGTAGCGTTGC |
| pHERD-*Pa*BphPS261A_fwd | GGCGTGCGCGCCTCGCTGGCGATATCCATCGTGGTCGGC |
| pHERD-*Pa*BphPS261A_rev | GCCGACCACGATGGATATCGCCATCGAGGCGCGCACGCC |
| pET21b-*At*BphP2D783N_fwd | GACGTCGCCATTCTCAACATCAATCTTGGATCCGACACC |
| pET21b-*At*BphP2D783N_rev | GGTGTCGGATCCAAGATTGATGTTGAGAATGGCGACGTC |
| pET21b-*Av*BphP2D793A_fwd | ACAGTTCCTGCCGTGGCAGTACTCGCCATCAACCTTGGCAATCATACC |
| pET21b-*Av*BphP2D793A_rev | GGTATGATTGCCAAGGTTGATGGCGAGTACTGCCACGGCAGGAACTGT |
